# Redefining Pleiotropic Drug Resistance in a Pathogenic Yeast: Pdr1 Functions as a Sensor of Cellular Stresses in *Candida glabrata*

**DOI:** 10.1101/2023.05.07.539747

**Authors:** Andrew N. Gale, Matthew W. Pavesic, Timothy J. Nickels, Zhuwei Xu, Brendan P. Cormack, Kyle W. Cunningham

## Abstract

*Candida glabrata* is a prominent opportunistic fungal pathogen of humans. The increasing incidence of *C. glabrata* infections is attributed to both innate and acquired resistance to antifungals. Previous studies suggest the transcription factor Pdr1 and several target genes encoding ABC transporters are critical elements of pleiotropic defense against azoles and other antifungals. This study utilizes *Hermes* transposon insertion profiling to investigate Pdr1-independent and Pdr1-dependent mechanisms that alter susceptibility to the frontline antifungal fluconazole. Several new genes were found to alter fluconazole susceptibility independent of Pdr1 (*CYB5*, *SSK1*, *SSK2*, *HOG1*, *TRP1*). A bZIP transcription repressor of mitochondrial function (*CIN5*) positively regulated Pdr1 while hundreds of genes encoding mitochondrial proteins were confirmed as negative regulators of Pdr1. The antibiotic oligomycin activated Pdr1 and antagonized fluconazole efficacy likely by interfering with mitochondrial processes in *C. glabrata*. Unexpectedly, disruption of many 60S ribosomal proteins also activated Pdr1, thus mimicking the effects of the mRNA translation inhibitors. Cycloheximide failed to fully activate Pdr1 in a cycloheximide-resistant Rpl28-Q38E mutant. Similarly, fluconazole failed to fully activate Pdr1 in a strain expressing a low-affinity variant of Erg11. Fluconazole activated Pdr1 with very slow kinetics that correlated with the delayed onset of cellular stress. These findings are inconsistent with the idea that Pdr1 directly senses xenobiotics and support an alternative hypothesis where Pdr1 senses cellular stresses that arise only after engagement of xenobiotics with their targets.

**Importance:** *Candida glabrata* is an opportunistic pathogenic yeast that causes discomfort and death. Its incidence has been increasing because of natural defenses to our common antifungal medications. This study explores the entire genome for impacts on resistance to fluconazole. We find several new and unexpected genes can impact susceptibility to fluconazole. Several antibiotics can also alter the efficacy of fluconazole. Most importantly, we find that Pdr1 – a key determinant of fluconazole resistance – is not regulated directly through binding of fluconazole and instead is regulated indirectly by sensing the cellular stresses caused by fluconazole blockage of sterol biosynthesis. This new understanding of drug resistance mechanisms could improve the outcomes of current antifungals and accelerate the development of novel therapeutics.

## INTRODUCTION

Candidiasis and other fungal infections impact millions of people throughout the world and cause thousands of deaths every year. *Candida albicans* is currently the most common cause of candidiasis with *C. glabrata*, a distantly related yeast, as the second most common. In the USA, the percentage of bloodstream infections (candidemia) due to *C. glabrata* rose from 12% to 29% between 1999 and 2012 (1) and to 38% in 2020, even surpassing *C. albicans* in that location (2). The steep rise of *C. glabrata* infection frequencies has been attributed to a combination of innate resistance to azole-class antifungals, easily acquired resistance to other antifungals, and nosocomial spread (3, 4). The CDC has classified such drug-resistant *Candida* species as serious threats.

The frontline antifungal fluconazole, as well as other azoles, targets lanosterol 14-alpha-demethylase (Erg11/Cyp51), a cytochrome P450 enzyme in the endoplasmic reticulum of fungal cells that is necessary for biosynthesis of ergosterol (5). Most *C. glabrata* clinical isolates are innately resistant to fluconazole due to two intrinsic regulatory pathways (6). The first pathway involves the transcription factor Upc2A, which drives expression of Erg11 and other enzymes of the ergosterol biosynthesis pathway upon ergosterol depletion (7). The second pathway involves the transcription factor Pdr1 and the inducible expression of target genes such as *CDR1*, *PDH1*, *SNQ2*, and *YOR1* genes, which all encode ABC-transporters that efflux a spectrum of compounds such as fluconazole, cycloheximide, and oligomycin (8–11). A prominent model proposes that direct binding of fluconazole, cycloheximide, and other xenobiotics to the core domain of Pdr1 (12) relieves auto-inhibition and promotes binding to PDRE-elements upstream of target genes (13) including *PDR1* itself (14). DNA-bound Pdr1 then recruits the mediator complex and RNA polymerase via a C-terminal activation domain (12, 15). Upc2A also participates in the second pathway by increasing *PDR1* and *CDR1* expression (16, 17). Overall, the model resembles that of the PXR transcription factors in humans, which confer pleiotropic drug resistance in tumors through increased expression of the ABC transporter P-glycoprotein/MDR1.

Mutations in Erg11 with reduced affinity for fluconazole confer drug resistance and therapeutic failures in patients infected with *C. albicans*, but such mutations arise very infrequently in *C. glabrata* (18). Instead, gain-of-function mutations in Pdr1 that appear to increase *CDR1* gene expression are often identified among hyper-resistant isolates of *C. glabrata* (19). Gain-of-function Pdr1 variants of *C. glabrata* also exhibited hyper-virulence in mouse models of invasive candidiasis even in the absence of antifungals (9, 20, 21). Thus, Pdr1 can promote virulence in addition to antifungal resistance. Alternatively, mitochondrial dysfunction acquired through mutations in the mitochondrial or nuclear genes results in constitutively high Pdr1 activity and *CDR1* expression, resulting in fluconazole resistance, enhanced virulence, and therapeutic failure (22–25). Transient mitochondrial dysfunction has been proposed as an adaptive strategy for *C. glabrata* survival and proliferation in fluconazole (24) and within phagosomes of macrophages (26). Precisely how mitochondrial dysfunction influences Pdr1 activity remains unclear. Nevertheless, similar phenomena may promote virulence and drug-resistance in *C. albicans* and other fungal species (27).

A better understanding of Pdr1 regulation in *C. glabrata* could improve therapeutic outcomes. To help clarify the cellular processes that regulate Pdr1 in *C. glabrata*, we present the results of genome-wide screens for genes influencing fluconazole resistance in *pdr1Δ* and *cdr1Δ* knockout mutants of BG14, a derivative of the vaginal isolate BG2 that exhibits innate resistance to fluconazole (28). This transposon-based approach (29) allows discrimination of genes that regulate fluconazole resistance independent of Pdr1 and Cdr1 such as the Upc2A-ERG pathway from those that depend on Pdr1 and Cdr1. These screens and follow-up experiments confirm that the Upc2A-ERG and Pdr1-Cdr1 pathways are the major determinants of innate fluconazole resistance in BG14 and operate independently. Surprisingly, we obtained evidence that Pdr1 is not activated by direct binding of fluconazole or other xenobiotics and propose that Pdr1 responds to stresses generated by deficiencies in Erg11, mRNA translation, and mitochondrial function. This updated model provides new opportunities for controlling pleiotropic drug resistance in *C. glabrata*.

## RESULTS

### Genes that increase fluconazole resistance independent of Pdr1 and/or Cdr1

*Hermes* transposon insertion profiling was utilized previously in *C. glabrata* strain BG14 to identify hundreds of genes that altered susceptibility to fluconazole (29). To determine which of these genes depend on Pdr1 and Cdr1, new pools of insertion mutants were generated in BG14 and isogenic *pdr1Δ* and *cdr1Δ* knockout mutants using an improved protocol (see Methods). The pools were cultured for 6.7 population doublings in synthetic complete dextrose medium containing or lacking 128 µg/mL fluconazole. The pools were then allowed to recover in drug-free medium and insertion sites were then amplified, sequenced, mapped to the BG2 reference genome (30), and tabulated gene-wise as before (29). Insertions within *UPC2A* were strongly under-represented in all three pools after exposure to fluconazole (Fig. 1), as expected because Upc2A induces *ERG11* gene expression and ergosterol biosynthesis independent of Pdr1 and Cdr1 (10). To estimate significance, z-scores were calculated (29) for every annotated gene (Sup. Tab. 1). The z-scores for *UPC2A* were strongly negative in the BG14, *pdr1Δ,* and *cdr1Δ* strains (−9.8, −4.2, and −9.5, respectively). These findings suggest the experimental conditions successfully detected genes that regulate fluconazole resistance independent of Pdr1 and Cdr1.

**Figure 1.**
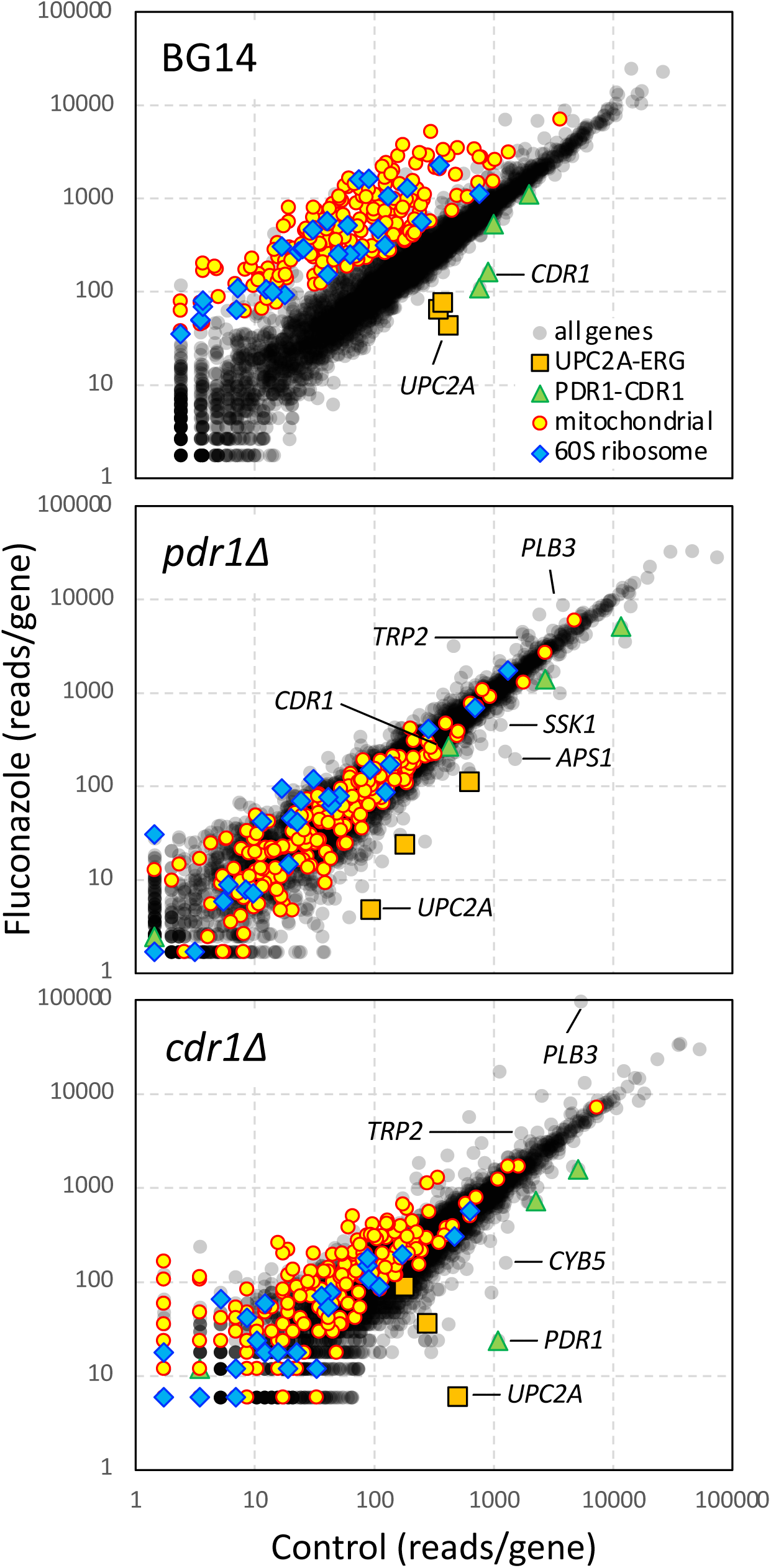
Genome-wide analyses of fluconazole susceptibility in *C. glabrata*. Diverse pools of *Hermes* transposon insertion mutants were generated in three isogenic derivatives of strain BG2 (BG14, *pdr1Δ*, *cdr1Δ*) and diluted into fresh medium containing or lacking 128 µg/mL fluconazole. The insertion sites were profiled using QI-seq, tabulated gene-wise, normalized, and plotted. Specific sets of genes were colorized: PDR1-CDR (green), UPC2A-ERG (orange), mitochondria (yellow), 60S ribosome (blue). Other genes are specifically labeled.

Resistance to fluconazole was also diminished when *DAP1* and *YHR045w* were disrupted with transposons in all three strain backgrounds, consistent with the earlier hypothesis that these mutants have diminished Erg11 function (29). The z-scores were strongly negative for *DAP1* (−6.8, −6.7, −1.2) and *YHR045w* (−6.8, −4.2, −3.7) in the BG14, *pdr1Δ*, and *cdr1Δ* strains, respectively. Only two other genes (*RIM15*, *MID2*) exhibited z-scores less than −3.0 in all three strains. Insertions in five additional genes (*APS1, YGK3, CYC8, DOA1, MPT5*) produced significantly negative z-scores in both the *cdr1Δ* and *pdr1Δ* strains but not the BG14 parent strain (Sup. Table 1). These genes are not known to function in any common pathways in *S. cerevisiae* and their possible contributions to ergosterol biosynthesis or alternative fluconazole resistance pathway is not known. An *aps1Δ* knockout mutation was introduced into all three strains and fluconazole susceptibility was assessed directly in broth microdilution assays. An *aps1Δ* knockout mutation was introduced into all three strain backgrounds and the concentration of fluconazole causing 50% growth inhibition (IC50) in broth microdilution assays was calculated. Consistent with the transposon datasets, the *aps1Δ* knockout mutation significantly diminished fluconazole resistance in the *pdr1Δ* and *cdr1Δ* backgrounds but not in the BG14 parent strain (Table 1).

**Table 1.** Fluconazole IC50’s of single and double knockout mutants.

|  | **BG14** | ***pdr1Δ*** | ***cdr1Δ*** |  |
| --- | --- | --- | --- | --- |
| **fluconazole** | 83 | 5.8 | 4.8 | IC50 (μg/mL) |
| ***aps1Δ*** | -1.5 | -1.5 | -1.7 | IC50 (fold change) |
| ***cyb5Δ*** | -3.6 | -4.6 | -2.9 |  |
| ***ssk1Δ*** | -1.0 | -1.9 | -1.0 |  |
| ***trp1Δ*** | 2.2 | 2.0 | 1.9 |  |
| ***cin5Δ*** | -1.7 | -1.1 | -1.0 |  |
| ***rpl9aΔ*** | 3.4 | 1.0 | 1.3 |  |

Insertions within *CDR1* strongly decreased resistance to fluconazole in the wild-type strain but not in the *pdr1Δ* strain (z-scores = −11.0, −1.7) as expected because *CDR1* is not expressed in *pdr1Δ* strains (9, 10). On the other hand, insertions in *PDR1* strongly decreased fluconazole resistance in both wild-type and *cdr1Δ* strains (z-scores = −11.3, −10.2). This finding is consistent with previous studies (8, 10) showing that Pdr1 can induce expression of *PDH1* and *SNQ2*, which independently confer weak resistance to fluconazole in *cdr1Δ* strains (11, 31). However, *PDH1*, *SNQ2*, and all other targets of Pdr1 (10, 13, 32) individually did not contribute significantly to fluconazole resistance even when disrupted in the *cdr1Δ* strain, indicating little individual impact of those genes in the conditions employed here.

Insertions in 13 genes significantly diminished resistance to fluconazole in *cdr1Δ* mutants but not *pdr1Δ* mutants or BG14. Within this set, 4 genes play roles in ER-associated protein degradation (*MNL1, HRD1, USA1, UBR1*) while 2 additional genes (*CYB5, TSC10*) participate in ER-associated biosynthesis of lipids (Sup. Table 1). Surprisingly, *cyb5Δ* knockout mutations decreased the IC50 of fluconazole in all three backgrounds instead of only one (Table 1), and therefore *CYB5* behaved similarly to *UPC2A*, *DAP1*, and *YHR045W*. In *S. cerevisiae*, *CYB5* encodes cytochrome b5, an ER-localized protein that enhances Erg3 and Erg5 activities but also enhances Erg11 function (33). The failure to detect *CYB5* as significant in two of the three pools may reflect the much lower density of transposon insertions recovered in this gene in those pools. Alternatively, the discordant behaviors of knockout mutants in isolation and in pools of transposon insertion mutants may reflect differences between the experimental conditions. To test this latter possibility, the *cyb5Δ* mutant and BG14 parent strain were mixed at 1:10 ratio and cultured in fluconazole conditions that mimic the transposon screen. In this configuration, *cyb5Δ* and wild-type BG14 strains grew at similar rates (Sup. Fig. 1), suggesting that the behavior of Cyb5-deficient mutants is dependent on the environmental conditions in which it is studied.

Another 25 genes conferred significant resistance to fluconazole in *pdr1Δ* mutants but not *cdr1Δ* mutants or BG14. Two of these genes (*IRA1, IRA2*) regulate the small GTPase RAS and are consistent with previous work that showed cAMP regulates fluconazole susceptibility in *S. cerevisiae* (34) and *C. albicans* (35). Two other genes (*DCS1, DCS2*) promote mRNA decay and other processes in *S. cerevisiae* (36). Additionally, 3 genes that function within the osmo-sensing branch of the HOG pathway (*SSK1, SSK2, HOG1*) (37) were part of this group. Introduction of a *ssk1Δ* mutation into all three strain backgrounds revealed a 1.89-fold decreased resistance to fluconazole in the *pdr1Δ* strain with no changes in fluconazole susceptibility to the *cdr1Δ* or BG14 strains (Table 1).

In summary, the genome-wide screens suggest that *C. glabrata* strain BG14 primarily utilizes the Pdr1-Cdr1 pathway and the Upc2A-ERG pathway in conjunction with Dap1, Yhr045w, Cyb5 for its innate resistance to fluconazole. Other genes and pathways such as *SSK1* conferred weaker fluconazole resistance through complex interactions with these or other pathways. While additional genes and pathways of fluconazole resistance may also operate in *C. glabrata*, their contributions in the BG2 background seemed less potent than the two major pathways.

### Genes that decrease fluconazole resistance independent of Pdr1 or Cdr1

Transposon disruption of 68 genes significantly increased resistance to fluconazole in *cdr1Δ*, *pdr1Δ*, or both strain backgrounds (Sup. Tab. 1), suggesting the normal gene decreases fluconazole resistance. Insertions in 4 genes (*TRP2, AVO1, INP52, CCW12*) increased fluconazole resistance in all three strain backgrounds. *TRP2* is one of five genes required for tryptophan biosynthesis in *S. cerevisiae*. Insertions in *TRP1*, *TRP3*, *TRP4*, and *TRP5* also conferred significant resistance to fluconazole in two of the three strains suggesting all five genes function similarly. To test whether tryptophan biosynthesis alters fluconazole susceptibility independent of the Pdr1-Cdr1 pathway, a *trp1Δ* knockout mutation was introduced into all three strains and tested for fluconazole susceptibility. The *trp1Δ* knockout mutation significantly increased fluconazole resistance in all three strain backgrounds (Table 1). A *trp1*Δ mutation in the CBS138 strain background *of C. glabrata* also increased fluconazole resistance by 2.0-fold (data not shown). Thus, tryptophan biosynthesis appeared to increase fluconazole susceptibility in *C. glabrata* independent of Pdr1 and Cdr1 through an unknown mechanism. These findings impact the interpretation of hundreds of gene knockouts that were analyzed for fluconazole susceptibility in a *trp1Δ* derivative of strain CBS138 (38).

Insertions in 7 genes (*MAK3, MAK10, PLB3, PTM1-B, GTB1, GPA2, PMR1-B*) significantly increased fluconazole resistance in both the *cdr1Δ* and *pdr1Δ* strains but not the BG14 control strain (Sup. Table 1). The products of *MAK3* and *MAK10* in *S. cerevisiae* bind each other and form the NatC-type N-terminal acetyltransferase that acylates the Golgi-associated protein Arl3, though *ARL3* did not appear significant in the screens. The products of *PTM1-B* and *PMR1-B* also are likely to function in the Golgi. Disruption of 24 genes increased fluconazole resistance in the *pdr1Δ* strain but not the *cdr1Δ* strain or control strain, with 5 of them involved in N-linked glycosylation in the ER (*ALG3, ALG5, ALG6, ALG8, CNE1*). Three additional genes in this group function in the ER and Golgi complex (*LAG1, AUR1-B, RER1*). Thus, insertions in these ER and Golgi genes exhibited the same strain dependences as insertions in *SSK1* and *SSK2* but with the opposite effect. Insertion mutations in another 29 genes increased fluconazole resistance in *cdr1Δ* strain but not the *pdr1Δ* strain, 17 of which are involved in mitochondrial functions. These mitochondrial mutants likely activate Pdr1 (29) and induce expression of *PDH1* and *SNQ2* that confer weak fluconazole resistance in the *cdr1Δ* background.

In summary, the genome-wide transposon screens suggest that additional negative regulators of the Upc2A-ERG (or some other) pathway conferring resistance to fluconazole may operate in *C. glabrata,* including the TRP pathway, NatC complex, and a number Golgi and ER proteins. By identifying the genes and processes that alter fluconazole susceptibility independently of Pdr1 and Cdr1, the genes that modify activity of the Pdr1-Cdr1 pathway can be identified more precisely.

### Mitochondrial dysfunction promotes fluconazole resistance via Pdr1

Mitochondrial dysfunction has been shown previously to hyperactivate Pdr1 and to induce expression of Cdr1 in *C. glabrata*, causing strong fluconazole resistance (9, 13, 22, 24). In an earlier study, we identified 135 nuclear genes encoding mitochondrial proteins that conferred significant fluconazole resistance when disrupted by transposons (29). Here, with improved methods, disruption of 198 mitochondrial genes were found to confer significant resistance to fluconazole (Fig. 1). These genes participate in a wide array of functions including mtDNA replication, transcription, translation, protein import, complex assembly and maturation, TCA cycle, respiratory chain complexes, ATP synthase, lipid biosynthesis, membrane dynamics, and more. Importantly, none of the mitochondrial genes from the two studies (228 in total) significantly altered fluconazole susceptibility when disrupted in the *pdr1Δ* strain (Fig. 1). These findings suggest that mitochondrial dysfunction increases fluconazole resistance primarily through up-regulation of Pdr1 activity.

Next, we tested whether oligomycin can mimic mutants that are deficient in its target, mitochondrial ATP synthase. In checkerboard assays, low doses of oligomycin increased resistance of BG14 strains to fluconazole by 5.0-fold but had no such effect on the *pdr1Δ* mutant (Fig. 2). The antagonism index of oligomycin on fluconazole efficacy was reduced to 2.0 in the *cdr1Δ* mutant, confirming that Cdr1 is the major effector of Pdr1 while weaker effectors Pdh1 and Snq2 remain in this mutant (29). Importantly, an *atp1Δ* knockout mutant that lacks a critical subunit of ATP synthase exhibited strong resistance to fluconazole when compared to the BG14 parent strain and was not further antagonized by oligomycin (Fig. 2). Similar results were obtained using *atp1Δ* and *atp2Δ* mutants of the CBS138 background (Sup. Fig. 2). Oligomycin also failed to antagonize fluconazole efficacy on the *atp1Δ pdr1Δ* double knockout mutant, which exhibited low resistance to fluconazole similar to that of the *pdr1Δ* mutant (Fig. 2). Thus, oligomycin completely phenocopied the effects of transposon insertions in its mitochondrial target and potently antagonized fluconazole efficacy in *C. glabrata*.

**Figure 2.**
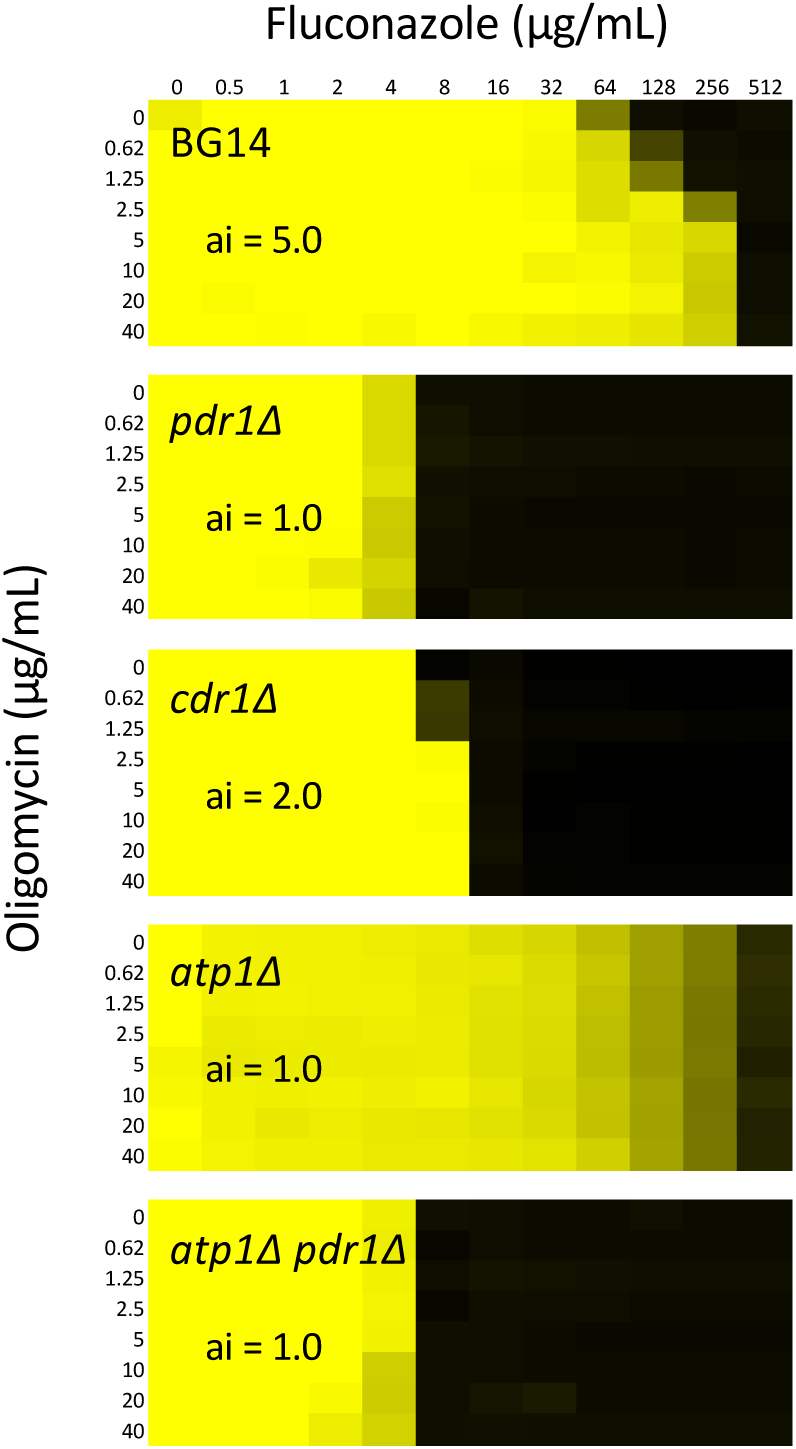
Mitochondria inhibitors antagonize fluconazole efficacy in Pdr1-dependent fashion. Growth of the wild-type BG14 and *pdr1Δ*, *cdr1Δ*, *atp1Δ*, and *atp1Δ pdr1Δ* derivative strains was measured at 600 nm after 20 hr incubation in SCD medium containing varying concentrations of fluconazole plus varying concentrations of either oligomycin. Data are plotted as a heatmap from saturation (yellow) to no growth (black). The IC50 of fluconazole was calculated for each row and the antagonism index (ai) was calculated for each dose of mitochondria inhibitor. The maximum ai for each strain and mitochondria inhibitor is indicated.

While hundreds of mitochondrial genes conferred Pdr1-dependent resistance to fluconazole when disrupted with transposons, insertions in six mitochondrial genes (*KGD1*, *KGD2*, *LSC1*, *LSC2*, *SDH9*, *SDH5*) significantly diminished fluconazole resistance in BG14 but not *pdr1Δ* or *cdr1Δ* strains (Sup. Table 1). These six genes are required for three successive steps of the TCA cycle in the mitochondrial matrix and inner membrane: conversion of alpha-ketoglutarate to succinyl-CoA, succinate, and fumarate. The z-scores of these six genes (−3.3, −3.1, −4.2, −2.9, −3.8, −5.8, respectively) in BG14 differ dramatically from those that perform the immediate upstream (*IDH1, IDH2*) and downstream (*FUM1*) reactions (+14.2, +9.1, and +10.7, respectively) of the TCA cycle. To test whether alpha-ketoglutarate, the product of *IDH1-IDH2* and the substrate of *KGD1-KGD2* gene products, might function as a negative regulator of Pdr1 or Cdr1, the *idh2Δ kgd2Δ* double knockout mutant was constructed and compared to the single mutants and the BG14 control strain. The *idh2Δ kgd2Δ* double mutant was highly resistant to fluconazole similar to the *idh2Δ* single mutant while the *kgd2Δ* single mutant remained less resistant than wild-type (Fig. 3A). Disruption of *PDR1* in *idh2Δ* and *kgd2Δ* strains nullified all their differences in fluconazole susceptibility (29). Therefore, mitochondrial biosynthesis of alpha-ketoglutarate appeared to be critical for decreased resistance of *kgd2Δ* mutants to fluconazole, suggesting that alpha-ketoglutarate, or its derivatives, may somehow down-regulate the activities of Pdr1 or Cdr1.

**Figure 3.**
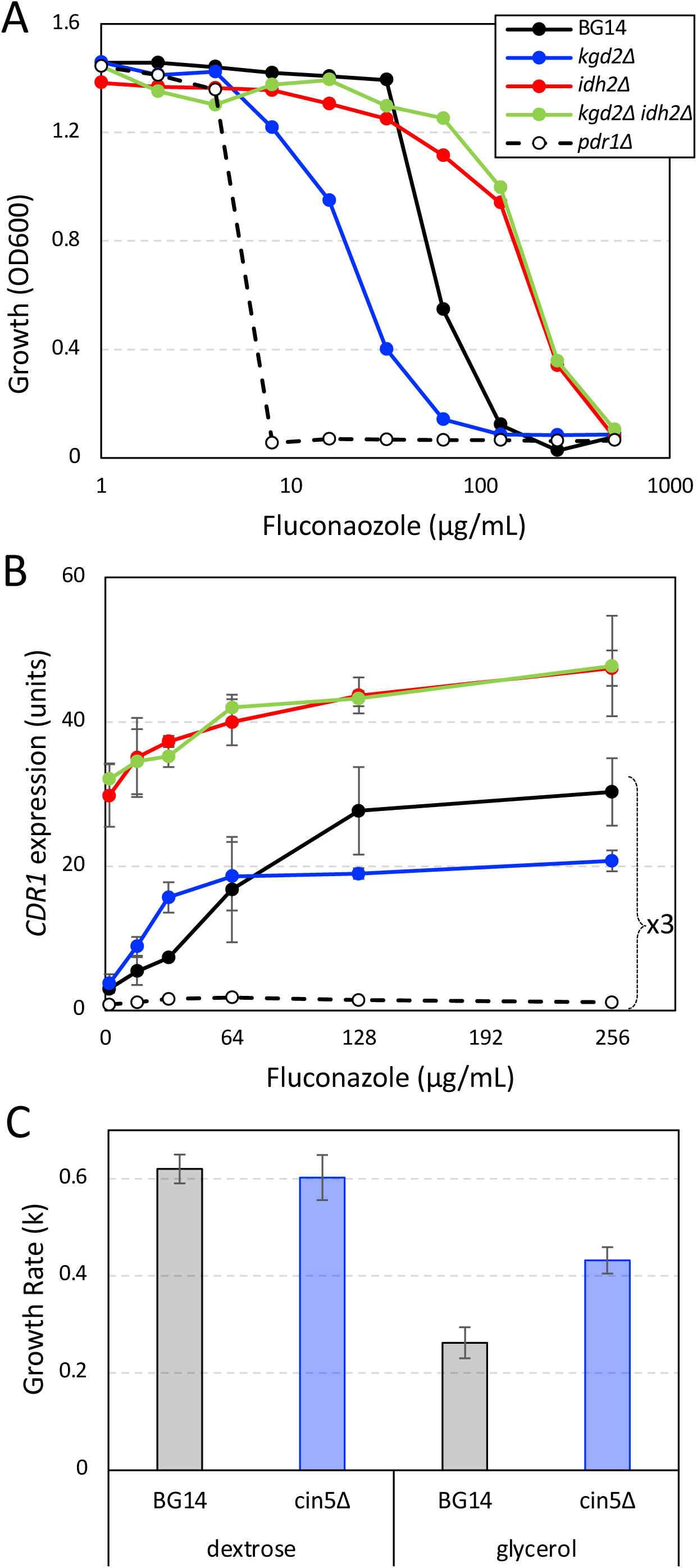
Mitochondrial dysfunction alters *CDR1* expression and fluconazole susceptibility. (A) Growth of *kgd2Δ idh2Δ* double and single gene knockout mutants was measured at 600 nm (OD600) after 20 hr incubation at 30°C in SCD medium containing different doses of fluconazole. (B) *CDR1* gene expression relative to *TEF1* gene expression was measured by RT-PCR after 6 hr exposure of log-phase cells to the indicated concentrations of fluconazole in SCD medium at 30°C. The data from three biological replicates were averaged (±SD) and normalized to BG14 parent strain in the absence of fluconazole. Values for BG14, *kgd2Δ*, and *pdr1Δ* were multiplied by 3 to improve visibility. (C) Three independent cultures of strains BG14 and *cin5Δ* were grown to log phase in synthetic complete medium containing 2% dextrose or 2% glycerol while periodically measuring optical density at 600 nm. Exponential growth rates (k = ln2/doubling time) were averaged and charted (±SD).

To help determine how alpha-ketoglutarate affects Pdr1 activity, *CDR1* and *TEF1* (control) mRNA levels were quantified by RT-PCR after exposure to varying concentrations of fluconazole for 6 hours. The *idh2Δ* and *idh2Δ kgd2Δ* both expressed high levels of *CDR1* mRNA relative to *TEF1* mRNA in the absence of added fluconazole relative to BG14 (defined as 1 unit in Fig. 3B) and both exhibited dose-dependent increases in *CDR1* expression (Fig. 3B). The *kgd2Δ* mutant strain and BG14 control strain expressed *CDR1* at low levels in the absence of fluconazole. Interestingly, the *kgd2Δ* mutant exhibited higher *CDR1* expression than BG14 at low doses and the opposite pattern at high doses (Fig. 3B). After fitting these data to sigmoid equations, maximal expression of *CDR1* was 1.6-fold lower in the *kgd2Δ* mutant relative to the BG14 control while half-maximal expression occurred at 3.1-fold lower concentration of fluconazole. These findings are consistent with an inhibitor of Pdr1 accumulating in the *kgd2Δ* mutant that lowers the maximal expression of *CDR1* and increases sensitivity to fluconazole. The *idh2Δ* and *idh2Δ kgd2Δ* strains may lack the inhibitory molecules or accumulate mitochondria-derived activators of Pdr1.

Insertion mutants that increase mitochondrial function may decrease resistance to fluconazole in wild-type strains but not in *pdr1Δ* and *cdr1Δ* backgrounds. A total of 75 genes were found to exhibit this pattern of fluconazole susceptibility in addition to the six genes of the TCA cycle described above (Sup. Table 1). One of the strongest candidate genes is *CIN5/YAP4* (z-score = −7.8 in BG14), encoding a bZIP transcription factor of the yAP-1 family that represses some mitochondrial genes in *S. cerevisiae* (39). A *cin5Δ* knockout mutant grew faster than BG14 control strain on medium containing the non-fermentable carbon source glycerol instead of glucose (Fig. 3C), suggesting that Cin5 may normally repress mitochondrial activity in *C. glabrata*. The *cin5Δ* mutant also exhibited diminished resistance to fluconazole in BG14 but not in *pdr1Δ* and *cdr1Δ* strain backgrounds (Table 1). Interestingly, the *CIN5* gene is naturally interrupted with a premature termination codon in two different patient isolates of *C. glabrata*, CBS138 (E160-stop; (40)) and DSY562 (Y62-stop; (41)). The *CIN5* locus from CBS138 was found to be non-functional when reintroduced into CBS138 or a *cin5Δ* derivative of BG14 while the reintroduced *CIN5* locus of BG14 was found to be functional (Sup. Fig. 3). These natural truncations in *CIN5* could be beneficial by improving cell growth in glucose-poor environments.

### Translation dysfunction increases fluconazole resistance via Pdr1 and Cdr1

Transposon disruption of 131 non-mitochondrial genes also increased resistance to fluconazole (z > 3.0) in BG14 but not in *pdr1Δ* or *cdr1Δ* strains. Interestingly, 30 of these genes (23%), encode subunits of the 60S ribosome or its assembly factors (Fig. 1; Sup. Table 1). This enrichment was striking because ribosomal genes are often small, essential, and poorly represented in the pools of transposon insertion mutants. Closer inspection of insertion sites revealed that exposure to fluconazole typically selected for insertions throughout the coding sequences of genes encoding non-essential ribosomal proteins (e.g. *RPL9A*, *RPL9B*, *RPL24A*, *RPP1A*, *RPP1B*). For essential ribosomal genes such as *RPL30* (Fig. 4A) and *RPL16B*, fluconazole often selected for insertions within the 3’ coding sequences and in the 5’ untranslated region. *RPL16B* truncation by insertion of the Tn7 transposon was previously found to confer resistance to fluconazole (24). These insertion mutants likely diminish the expression or function of the 60S ribosomal subunit resulting in hypomorphic mRNA translation.

**Figure 4.**
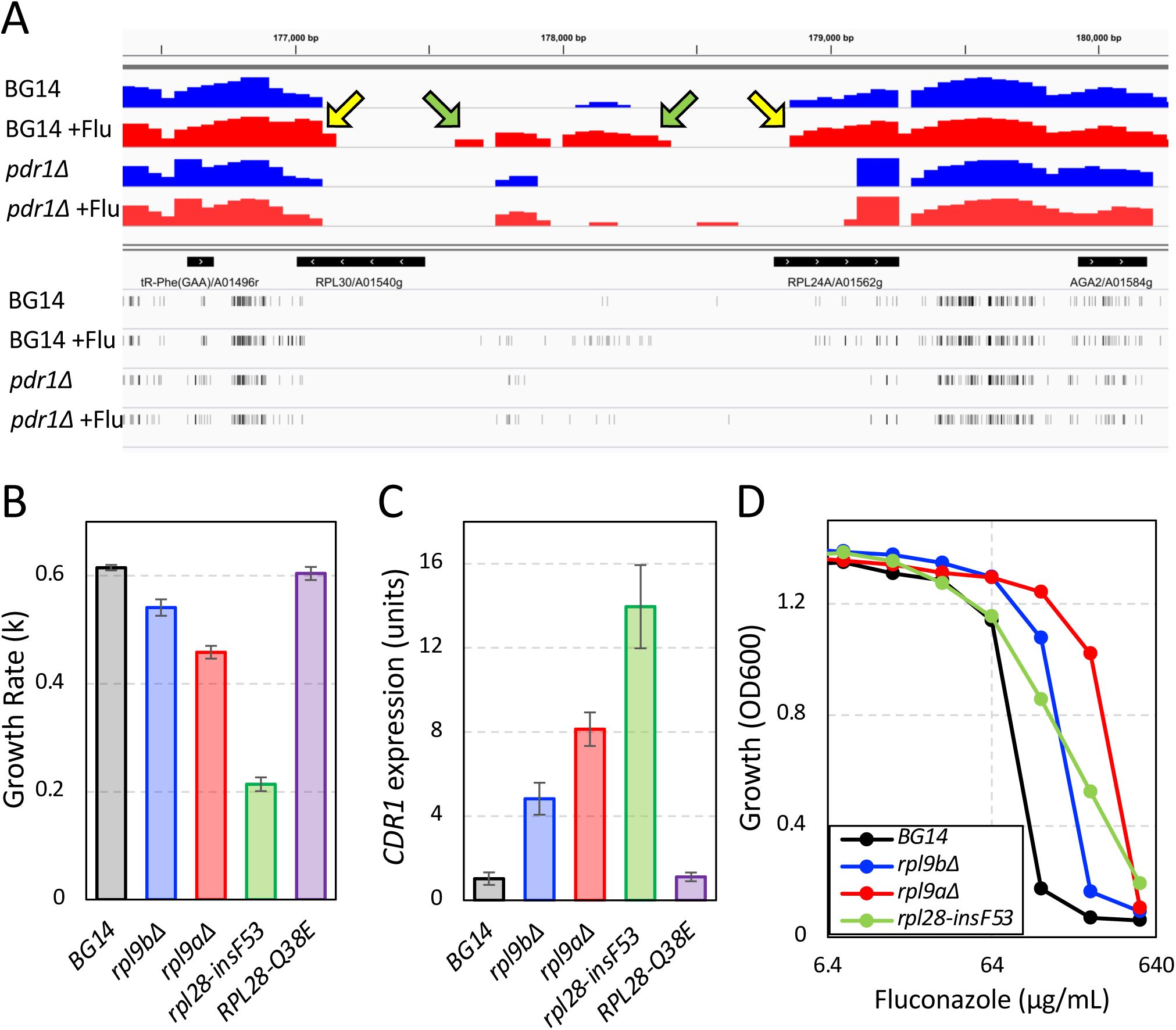
60S ribosome dysfunction alters *CDR1* expression and fluconazole susceptibility. (A) Transposon insertion sites and read counts were visualized in a 4 kilobase segment of chromosome A spanning divergently transcribed *RPL30* and *RPL24A* genes for the BG14 and *pdr1Δ* pools with (red) and without (blue) exposure to fluconazole. Histograms of read coverage were plotted on a log scale. Arrows indicate the 5’ noncoding regions (green) and the 3’ coding regions (yellow) that were enriched after fluconazole exposure in BG14 but not *pdr1Δ*. Exponential growth rates (B), basal *CDR1* expression levels (C), and fluconazole resistances (D) of several mutants of the 60S ribosome were quantified in SCD medium at 30°C as described in Methods.

To validate these findings, *rpl9aΔ* and *rpl9bΔ* knockout mutants and a 1-codon insertion mutant of *RPL28* (*rpl28-insF53*) were introduced into BG14 and studied. All three mutants proliferated at slower rates than the control strain (Fig. 4B) suggesting all were hypomorphic for mRNA translation. All these mutants exhibited elevated expression of *CDR1* in the absence of fluconazole (Fig. 4C) as well as strong resistance to fluconazole in growth assays (Fig. 4D). Therefore, genetic deficiencies in mRNA translation appeared to activate Pdr1 and induce Cdr1, resulting in enhanced fluconazole resistance.

### Cycloheximide activates Pdr1 indirectly via engagement of its target

Cycloheximide, a potent inhibitor of eukaryotic mRNA translation, has been shown to activate Pdr1 and induce *CDR1* expression in *C. glabrata* (9). Thakur et al. observed direct binding of cycloheximide to Pdr1 *in vitro* and proposed Pdr1 functioned as a direct xenobiotic receptor in cells (12). This hypothesis predicts that mutations that weaken cycloheximide binding to ribosomes will have no impact on Pdr1 activation by cycloheximide. To test this prediction a derivative of BG14 expressing an *RPL28-Q38E* variant was generated. The same mutation in *S. cerevisiae* diminished affinity of ribosomes for cycloheximide by about 10-fold without altering general translation rates (42). The *RPL28-Q38E* mutant of *C. glabrata* exhibited 8-fold increase in resistance to cycloheximide and no change in resistance to anisomycin, a distinct translation inhibitor that binds elsewhere on translating ribosomes (43) (Fig. 5A). After exposure to low doses of cycloheximide, the *RPL28-Q38E* mutant exhibited much less expression of *CDR1* than wild-type BG14 (Fig. 5B). In the absence of cycloheximide, the *RPL28-Q38E* strain grew at wild-type rates (Fig. 4B) and expressed *CDR1* at wild-type basal levels (Fig. 4C), suggesting that this mutant does not cause detectable translation distress. These findings are inconsistent with predictions of the xenobiotic receptor hypothesis and suggest that cycloheximide activates Pdr1 indirectly by first engaging ribosomes.

**Figure 5.**
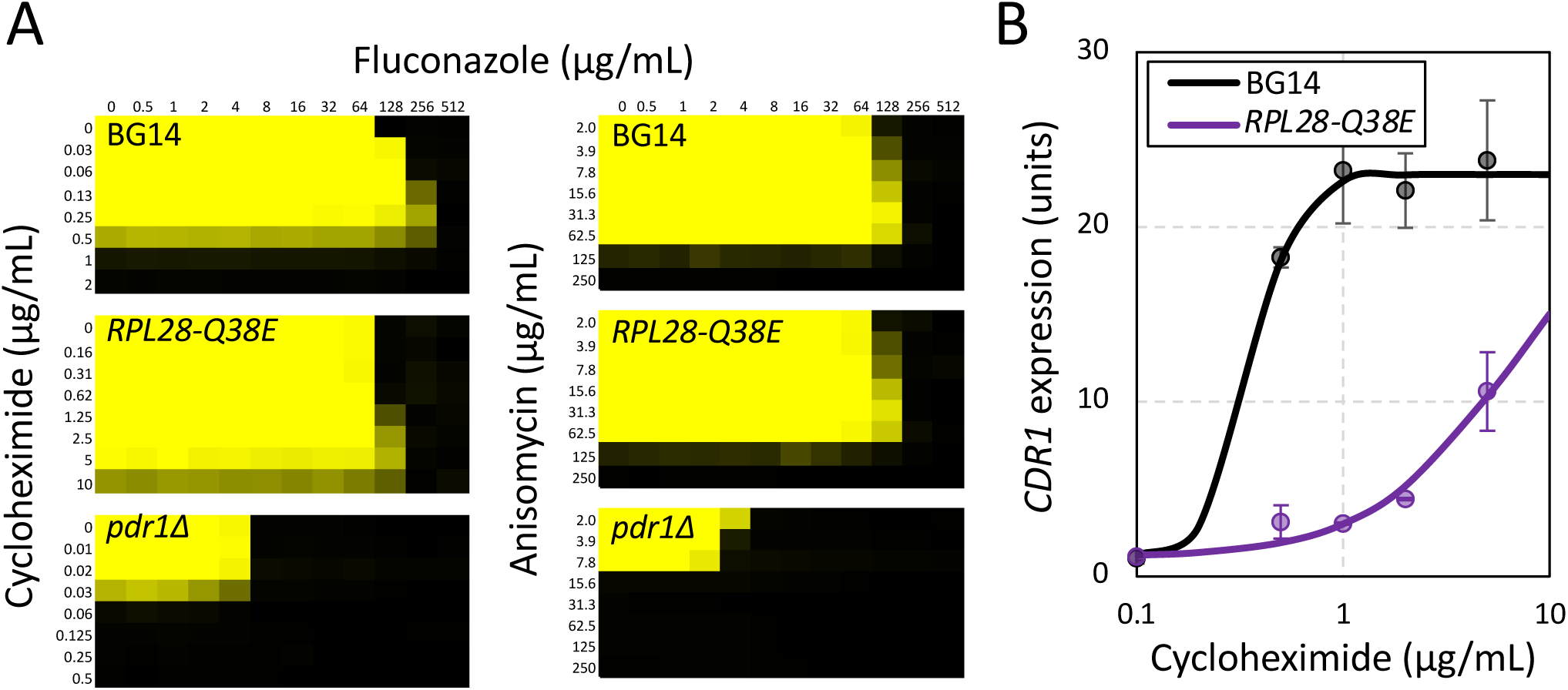
Translation inhibitors activate Pdr1 and antagonize fluconazole efficacy through engagement of ribosomes. (A) Checkerboard assays of BG14, *RPL28-Q38E*, and *pdr1Δ* mutants were determined using varying concentrations of fluconazole plus varying concentrations of cycloheximide (left) or anisomycin (right) as described in Fig. 2. (B) Expression of *CDR1* in wild-type (BG14) and *RPL28-Q38E* mutant strains of *C. glabrata* was measure by RT-PCR after 1 hr exposure to the indicated concentrations of cycloheximide in SCD medium at 30°C. Data points indicate the average (±SD) of three biological replicates and smooth curves represent best-fit sigmoid equations generated by non-linear regression.

Sub-lethal doses of cycloheximide have been found to antagonize the efficacy of fluconazole in *C. krusei* (44). Cycloheximide and anisomycin also antagonized fluconazole efficacy in *C. glabrata* strain BG14 but not in the *pdr1Δ* mutant (Fig. 5B) or *cdr1Δ* mutant (not shown). This finding suggests the translation inhibitors antagonize fluconazole by activating Pdr1 and inducing *CDR1* expression. If these inhibitors activate Pdr1 indirectly via inhibition of translation, cycloheximide antagonism should be diminished in the *RPL28-Q38E* mutant while anisomycin antagonism should remain similar in BG14. Indeed, this expectation was observed (Fig. 5B). Altogether, the findings argue against the hypothesis of Pdr1 as a direct sensor of cycloheximide. Instead, they suggest that Pdr1 senses deficiencies in mRNA translation resulting from translation inhibitors or from hypomorphic mutations in the 60S subunit.

### Fluconazole activates Pdr1 indirectly via engagement of its target and generation of cellular stresses

Purified Pdr1 binds fluconazole *in vitro* (12) but the impact of this binding has not been tested in *C. glabrata* cells. If Pdr1 directly senses fluconazole *in vivo*, overexpression of Erg11 or a double mutant variant termed Erg11-DM (45) that likely diminishes its affinity for fluconazole because of the Y141H substitution (46) would be expected to have little or no impact on Pdr1 activation by fluconazole. To test this prediction, plasmids bearing *ERG11* or *ERG11-DM* coding sequences under control of a strong constitutive promoter (47) were introduced into BG14 and then *CDR1* gene expression was quantified by RT-PCR after 6 hr growth in varying doses of fluconazole. Western blot experiments confirmed Erg11 and Erg11-DM proteins were overexpressed to similar degrees (Fig. 6A). Relative to the control strain, the Erg11 and Erg11-DM overexpressing strains exhibited 3.0-fold and 17-fold shifts in ED50, the dose of fluconazole that produced half-maximal expression of *CDR1* (Fig. 6B). Growth assays showed IC50s were shifted similarly (3.4-fold and 13-fold; Fig. 6C). Moreover, the same patterns were observed in a *upc2aΔ* knockout mutant (Sup. Fig. 4), though this strain was 16-fold less resistant to fluconazole than the BG14 parent and maximal *CDR1* expression was 1.3-fold lower. When integrated into its genomic locus, Erg11-DM was severely hypomorphic and conferred both slow growth and high constitutive expression of *CDR1* (47). Other slow-growing hypomorphs of Erg11 also activated Pdr1 and induced *CDR1* expression in the absence of any inhibitors (48). Altogether, these findings suggest that Erg11 deficiency is sufficient for Pdr1 activation, and that fluconazole must effectively inhibit Erg11 in order to activate Pdr1.

**Figure 6.**
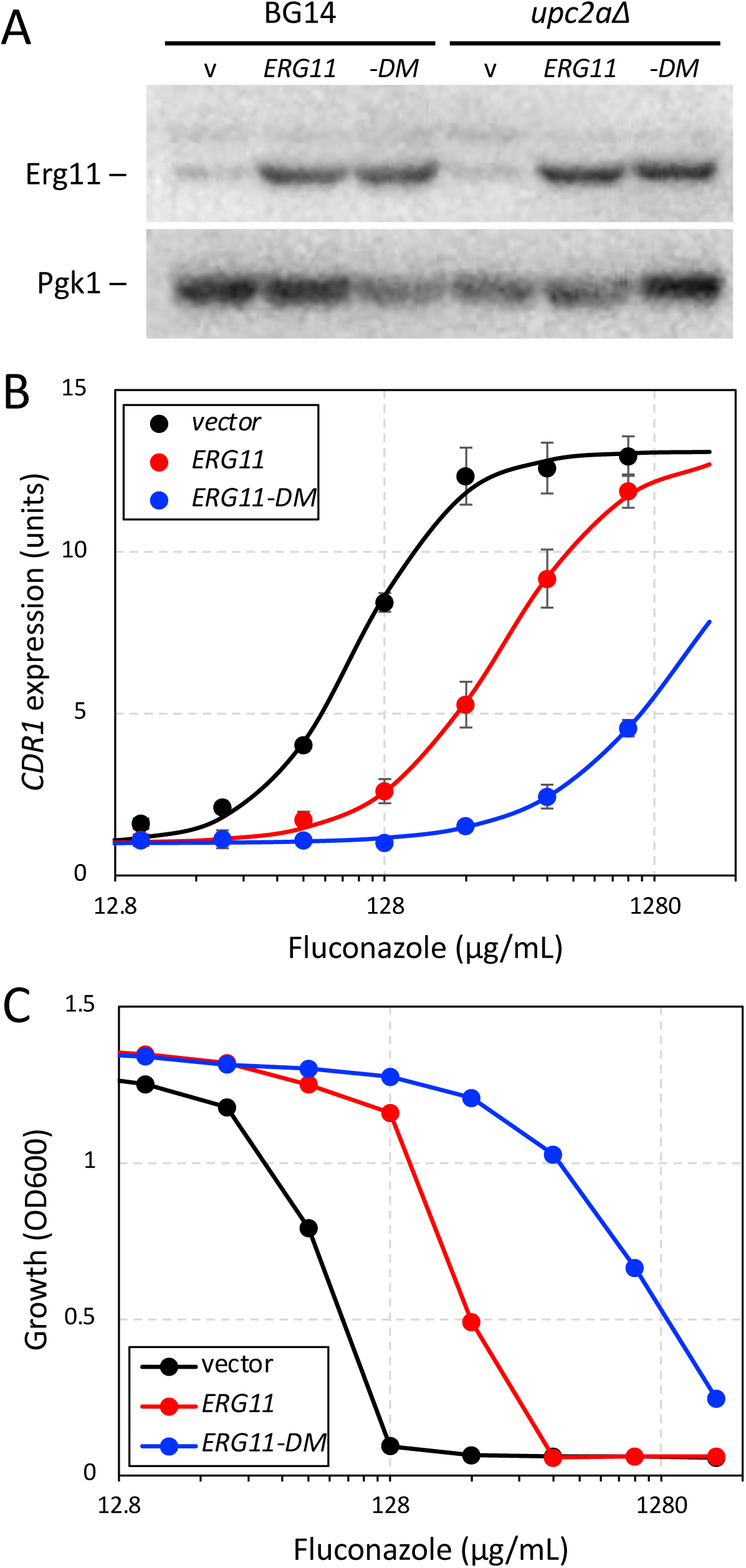
Fluconazole activates Pdr1 through engagement of Erg11. (A) Western blot analysis of Erg11 and Pdc1 proteins in lysates of BG14 and *upc2aΔ* cells bearing plasmids pCU-PDC1 (v), pCU-PDC1-ERG11 (ERG11), or pCU-PDC1-ERG11-Y141H,S410F (-DM). (B) The same strains were exposed to varying concentrations of fluconazole for 6 hr in SCD medium at 30°C, and then *CDR1* expression was quantified using RT-PCR as described in Fig. 3. Data points represent averages of 3 biological replicates (±SD) and smooth curves represent best-fit sigmoid equations generated by non-linear regression. (C) Fluconazole susceptibility of the same strains was determined as described in Fig 3.

If Pdr1 were to directly bind and sense fluconazole, cycloheximide, and other xenobiotics, the induction of *CDR1* expression would likely be rapid and consistent from compound to compound if cellular uptake is similar. To test these predictions, we measured *CDR1* expression in BG14 cells at different times following exposure to high doses of cycloheximide and fluconazole. *CDR1* expression was induced much faster in response to cycloheximide than to fluconazole (Fig. 7A). Ketoconazole, another azole-class inhibitor of Erg11, induced *CDR1* expression with slow kinetics like fluconazole while oligomycin induced *CDR1* expression with intermediate kinetics (Fig. 7A). Thus, there was little consistency in the kinetics of Pdr1 activation by these xenobiotics. The slow responsiveness to fluconazole and ketoconazole correlated with the delayed onset of growth inhibition, which first appeared after 90 minutes of drug exposure (Fig. 7B). The rate of Erg11 inhibition in these conditions has not been studied, so the possibility that azoles are taken up slowly has not been excluded. Nevertheless, Pdr1 activation kinetically correlated with the onset of cellular stresses generated by Erg11 inhibitors rather than the presence of the xenobiotics themselves.

**Figure 7.**
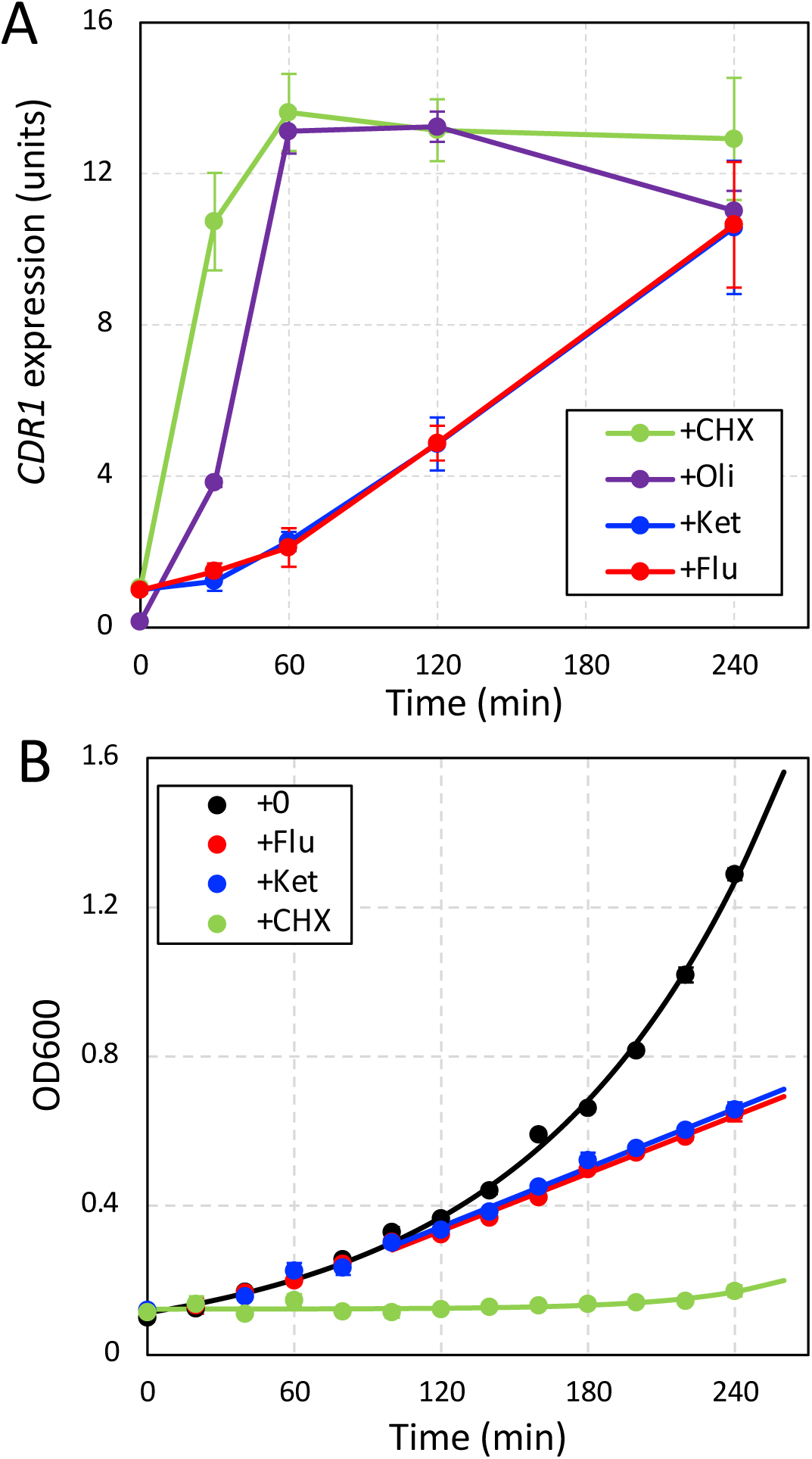
Kinetic analyses of Pdr1 activation and growth inhibition following exposure to cycloheximide and fluconazole. (A) *CDR1* expression was monitored by RT-PCR in BG14 cells at different times after exposure to 2 µg/mL cycloheximide, 40 µg/mL oligomycin, 256 µg/mL fluconazole, or 48 µg/mL ketoconazole. Data points represent averages (±SD) of 3 biological replicates. (B) Growth of BG14 in SCD medium at 30°C was monitored at 600 nm at different times after diluting log-phase cells into fresh medium containing cycloheximide, fluconazole, or ketoconazole at concentrations in A. Data points represent averages (±SD) of three biological replicates and smooth curves represent best-fit exponential (black, green) or linear (blue, red; t > 90 min) equations using regression.

In summary, fluconazole, cycloheximide, and oligomycin all activated Pdr1 indirectly via engagement of their established targets while genetic depletion of those targets activated Pdr1 independent of the xenobiotics. These findings are not consistent with the hypothesis that Pdr1 functions as a direct sensor of xenobiotics *in vivo* and suggest instead that Pdr1 somehow senses the cellular stresses that arise through dysfunction of ergosterol biosynthesis, mRNA translation, and mitochondria.

## DISCUSSION

This study makes use of forward genetic screens in *pdr1Δ* and *cdr1Δ* mutants of *C. glabrata* to establish several new insights into Pdr1-independent and Pdr1-dependent mechanisms of fluconazole resistance. The most notable findings argue against an earlier hypothesis that Pdr1 functions as a direct receptor of fluconazole and other xenobiotics. Instead, the findings support an alternative hypothesis where Pdr1 indirectly senses diverse cellular stresses that arise from the inhibitory action of xenobiotics on their established targets.

Key evidence for the indirect mechanism of Pdr1 activation derives from analysis of Pdr1 activity in strains where xenobiotic targets were mutated to have reduced affinity for their inhibitors. Low-affinity variants of Rpl28 and Erg11 strongly diminished the responsiveness of Pdr1 to cycloheximide and fluconazole, respectively. Overexpression of wild-type Erg11 diminished sensitivity of Pdr1 while underexpression of Erg11 and other enzymes in the ergosterol biosynthesis pathway (in strains lacking Upc2A) increased sensitivity of Pdr1 to fluconazole. Additionally, hypomorphic mutations in Erg11 (48) and 60S ribosomes (Fig. 4) activated Pdr1 in the absence of any xenobiotics. Finally, unlike cycloheximide, fluconazole and ketoconazole activated Pdr1 with very slow kinetics that correlated with a delayed onset of cellular stress (growth rate inhibition). Though we cannot rule out the possibility that slow cellular uptake of the Erg11 inhibitors delays their effects, cellular stresses appear to be necessary and sufficient for Pdr1 activation while the xenobiotics themselves were neither. All these findings are inconsistent with predictions of the xenobiotic receptor hypothesis, where Pdr1 was proposed as a direct sensor of cycloheximide and fluconazole (12).

Another concern with the xenobiotic receptor hypothesis arises from reports of the very low affinity of *S. cerevisiae* Pdr1 for ketoconazole (Kd = 39 µM) as compared to the 3000-fold higher binding affinity of ketoconazole to Erg11 from *C. albicans* with (Kd = 12 nM) (12, 49). If a similar disparity were to exist for the *C. glabrata* proteins, moderate ketoconazole concentrations would strongly inhibit Erg11 function while Pdr1 would remain almost completely unbound and therefore ineffective in promoting expression of *CDR1*. Variants of Pdr1 with diminished affinity for xenobiotics are predicted by the receptor hypothesis to be deficient in *CDR1* expression in response to the compounds. However, no such variants have been reported and deletion or mutation of the xenobiotic-binding core domain of Pdr1 often results in high constitutive expression of *CDR1* (14). The significance of xenobiotic binding to Pdr1 therefore remains unresolved *in vivo* and likely to be secondary to xenobiotic action on the primary target. More detailed studies will be required to clarify the biological impacts of xenobiotic binding to Pdr1.

Mitochondrial dysfunction caused by either mutations or inhibitors also activated Pdr1, likely through multiple interacting mechanisms (27). One of these mechanisms may include the negative regulation of Pdr1 by mitochondria-synthesized alpha-ketoglutarate, or one of its derivatives. In *C. glabrata*, *kgd2Δ* mutants that fail to consume mitochondrial alpha-ketoglutarate exhibited low Pdr1 activity and diminished resistance to fluconazole. In contrast, the *idh2Δ kgd2Δ* double mutants that synthesize little mitochondrial alpha-ketoglutarate exhibited high constitutive Pdr1 activation and resistance to fluconazole even though the double mutants are as deficient in respiration as *kgd2Δ* mutants. Alpha-ketoglutarate deficiency may help explain the fluconazole-resistant phenotype of hundreds of other mitochondria-deficient mutants, some of which have arisen naturally in patients with fluconazole-resistant infections (23). Alternatively, isocitrate (or its derivatives) that is expected to increase in *idh2Δ* mutants could somehow increase Pdr1 activity. AlphaFold predicts at least one deep pocket in the core domain of Pdr1 that could potentially bind small metabolites (https://alphafold.ebi.ac.uk/entry/B9VI15). While the function of this hypothetical deep pocket remains untested, this study provides a framework for investigating how Pdr1 senses diverse cellular stresses and thereby achieves multidrug resistance.

At present, the mechanisms by which mitochondrial dysfunction, ergosterol depletion, and translational deficiencies activate Pdr1 are not defined. The mechanisms may operate independently of one another or converge on a single regulatory network. Other compounds that activate Pdr1 and antagonize fluconazole may do so by generating mitochondrial, ergosterol, or translation stresses. The antifungal drug flucytosine activates Pdr1 and induces *CDR1* and fluconazole resistance in *C. glabrata* probably by interfering with mitochondrial function (50). The antimalarial drugs artemisinin and artesunate activated Pdr1 in *C. glabrata* (51) and *S. cerevisiae* (52) but also interfered with mitochondrial activities. Secondarily, the artemisinin-related molecules appeared to inhibit Cdr1 activity and therefore antagonism of fluconazole efficacy was not observed (53, 54). A screen of 1,280 FDA-approved drugs in *C. glabrata* revealed several additional compounds with unknown targets that antagonized fluconazole in a Pdr1-dependent fashion (55). A better understanding of Pdr1 regulatory mechanisms will shed light on the medications that antagonize and synergize with fluconazole.

*C. albicans* lacks Pdr1 and instead utilizes a distantly related transcription factor Tac1 to drive expression of *CDR1* and other efflux pumps (56). A different subset of compounds antagonized fluconazole in *C. albicans* by a Tac1-dependent process (55), suggesting this sensor may detect a different range of stresses than Pdr1 in *C. glabrata*. Several dozen hypomorphic mitochondrial mutations of *C. albicans* were recently found to exhibit resistance to fluconazole through enhanced activity of Cdr1 that was independent of Tac1 or other effects on expression (57). A mutant of *C. albicans* deficient in complex III of the respiratory chain increased resistance to fluconazole (58) while a mutant deficient in complex IV decreased resistance to fluconazole (59). Lastly, the experimental antifungal ML316, which inhibits a mitochondrial transporter of inorganic phosphate, strongly antagonized fluconazole efficacy in *C. albicans* (59). Because mitochondrial dysfunction is generally more lethal in petite-negative *C. albicans* relative to petite-positive *C. glabrata* or *S. cerevisiae*, there may be significant differences in the way fluconazole efflux pumps are regulated. While differences among Pdr1/Tac1-superfamily of transcription factors are likely in the different yeast species, some of the regulatory principles may be conserved and exploited for development of improved antifungal therapies.

This study also sheds light on mechanisms of fluconazole resistance that operate independent of Pdr1 in *C. glabrata*. In *pdr1Δ* mutants, *UPC2A* strongly conferred resistance to fluconazole, likely through its ability to induce expression of *ERG11* and other components of the ergosterol biosynthetic pathway (16) including *CYB5* (60). *CYB5* knockout mutations in *pdr1Δ* and control strains diminished resistance to fluconazole, likely due to diminished ergosterol biosynthesis as observed previously in *C. albicans* (61). Cyb5 is thought to supply electrons to Erg3 (62) and also to Erg11 and Erg5 when an alternative electron carrier Ncp1 was inactivated (33). However, *erg3-* insertion mutants and *erg3Δ* mutants of *C. glabrata* were not resistant to fluconazole (63). *erg3-* mutants arose spontaneously during prolonged passaging in anidulafungin but did not arise during *in vitro* evolution of fluconazole resistance where gain-of-function missense mutations in *PDR1* and *ERG11* predominated (64). The conditions employed in this study survey only loss-of-function mutations and their immediate responsiveness to high fluconazole, thereby providing a distinct perspective on acquired fluconazole resistance.

Different *C. glabrata* isolates often exhibit enormous genetic diversity that could adaptively alter many traits relevant to virulence, pathogenicity, and drug resistance (65). Findings presented above suggest the natural polymorphisms that truncate *CIN5* in isolates CBS138 and DSY562 may be adaptive and may also influence Pdr1 regulation and fluconazole susceptibility of those isolates. The natural truncation of *SSK2* in the isolate CBS138 alters susceptibility to osmotic stress (37) and may influence fluconazole susceptibility through post-transcriptional effects on Cdr1 function. A complete understanding of antifungal resistance mechanisms in *C. glabrata* will require parallel analyses in multiple different isolates and experimental conditions. The high portability of *Hermes* transposon insertion profiling may accelerate progress toward that goal.

## METHODS

### Plasmids, strains, and culture conditions

The plasmid pCU-MET3-Hermes (29) was transformed into isogenic wild-type (BG14), *pdr1Δ* (CGM1094), and *cdr1Δ* (CGM1096) strains of *C. glabrata* (66) using the lithium acetate protocol (28). To induce transposition, single colonies were inoculated into 100 mL of synthetic complete 2% dextrose (SCD) medium lacking uracil, cysteine, and methionine, divided equally into 40 separate 16 × 150 mm glass culture tubes, and shaken for three days at 30° C. The cultures were then pooled, recovered in fresh medium, and depleted of non-transposed cells by resuspending the pelleted cells in 300 mL of fresh SCD media containing 1 mg/mL of 5-fluoroorotic acid (ZymoResearch) and 0.1 mg/mL of nourseothricin (Gold Biotechnology) and shaking overnight at 30°C. This step was repeated an additional time. Next, 60 mL of the culture was pelleted and resuspended in 600 mL of fresh SCD media containing 1 mg/mL of 5-fluoroorotic acid and 0.1 mg/mL of nourseothricin and grown shaking overnight at 30°C. Cells were then pelleted and resuspended in 60 mL of 15% glycerol mixed with SCD and frozen at −80°C for long-term storage in 10 mL aliquots. This modified protocol was found to minimize jackpot insertions while increasing overall complexity of the pools relative to previous methods (29).

To perform drug screening, aliquots of each pool were thawed, washed once in SCD medium, and diluted 10-fold into SCD medium and allowed to shake overnight at 30°C. These cultures were then diluted 100-fold into SCD medium with or without 128 µg/mL of fluconazole (Cayman Chemicals) and allowed to grow shaking overnight at 30°C. Cells were then pelleted and washed 1x in SCD medium and resuspended into fresh SCD medium for overnight growth, shaking at 30°C. Cells were then pelleted and resuspended in 30 mL of 15% glycerol and 1x SCD medium and frozen at −80°C for long-term storage in 5 mL aliquots before undergoing the QI-seq protocol (67).

Sup. Table 2 contains a complete list of *C. glabrata* strains used in this study. *RPL28* mutants were generated by growing BG14 to saturation overnight in YPD medium and plating onto YPD plates containing 5 µg/mL of cycloheximide (Sigma). Colonies were purified on YPD plates and screened by PCR-seq for mutations in *RPL28* using primers listed in Sup. Table 3.

To generate the *kgd2Δ idh2Δ* double mutant (strain AGY49), strain AGY15 was converted to *ura3-* (AGY47) by selecting for spontaneous mutants able to grow on SCD plates containing 1 mg/mL of 5-fluoroorotic acid. A PCR-product containing *idh2Δ::URA3* locus with ∼500bp homology arms was amplified from strain AGY07 and transformed into AGY47. Successful *idh2Δ* knockouts were identified by PCR using screening primers (Sup. Table 3).

To generate *trp1Δ* mutants, the *trp1Δ::URA3* locus of CBS138-2001T (68) with ∼500 bp homology arms was amplified by PCR and transformed into strains BG14, CGM1094, and CGM1096. Colonies were purified and screened using PCR as well as inability to grow on SCD plates lacking tryptophan.

The *ssk1Δ*::NATr locus with ∼500 bp flanking homology was PCR amplified from a CBS138-derived strain (38) and transformed into strains BG14, CGM1094, and CGM1096. Colonies appearing on YPD plates containing 0.1 mg/mL nourseothricin were purified on YPD plates and screened by PCR.

All other mutants were knocked out using the PRODIGE method (69) and the PCR primers listed in Sup. Table 3. Briefly the *ScURA3* coding sequence was amplified with 60bp extensions homologous to the 5’ and 3’ untranslated regions of the target gene. The resultant PCR product was transformed into the relevant parent strains using the standard lithium acetate method and plated onto SCD-Ura plates. Colonies were purified and validated using screening primers. To create plasmids overexpressing ERG11 or ERG11-DM, the centromeric plasmid pCU-PDC1 (47) was digested with XhoI and XmaI, ligated to similarly digested PCR products obtained from genomic DNA of strains BVGC14 and BVGC340 (45), respectively, and transformed into DH5α *E. coli* cells. To create plasmids containing *CIN5*, pCN-PDC1 was digested with SacI and SpeI and ligated to PCR products obtained from CBS138 and BG14 gDNA templates from −500 to +1289 relative to the start codon. Plasmids were authenticated by DNA sequencing.

### Genomic extraction, QIseq, Data analysis

Genomic DNA was extracted using the Quick-DNA Fungal/Bacterial Miniprep kit (Zymo Research). Approximately of 100 mg of wet cell pellets were processed by the kit after three washes with 1 mL of deionized H_2_0. A total of 4 libraries with a starting material of 600 ng of gDNA were prepared for each condition using the QIseq method previously described (29). Briefly purified gDNA was sonicated using a Diagenode Picoruptor to an average size of 350 bp. The fragmented DNA was end repaired, ligated to indexed adapters, size selected using Ampure XP beads, and PCR amplified in two separate stages: the first using primers specific for the adapter and the transposon and the second using primers that added Illumina capture sequences to the fragments. Both PCR reactions were purified using AMPure XP beads. Libraries were sized using the Agilent Tape Station and replicate samples were pooled to equal molarity. Samples were sequenced on a MiSeq instrument (Illumina) with 2×76bp paired-end reads. Reads were demultiplexed using CutAdapt (70) and mapped to the BG2 reference genome (30) using Bowtie2 (71). Mapped reads were filtered for mapping quality and assigned to individual genes using a custom Python script. Bedfiles were generated and visualized using the IGV genome browser (72).

### Determination of IC50 and antagonism

To determine the IC50 of the strains to different drugs each strain was grown to saturation overnight in SCD medium. Saturated cells were diluted 1000-fold into fresh SCD medium and diluted another 2-fold into 75 µL SCD medium containing variable concentrations of the requisite drugs in a 96-well dish. The dishes were mixed, incubated for 20 hours at 30°C, mixed again, and optical density was measured at 600 nm (Accuris SmartReader 96T). IC50 was calculated from the raw data by non-linear regression using the sigmoid equation (KaleidaGraph v5). Antagonism was determined similarly, except that two drugs were varied simultaneously over broad concentration ranges on the same 96-well dish and IC50 was determined in rows and columns.

### Growth rate determination

To measure growth rates, strains were grown overnight at 30°C in SCD (2% glucose) or SCG (2% glycerol) medium to mid-log phase and then back diluted into fresh medium containing or lacking various drugs to an OD600 = 0.1. Three replicate cultures of each strain were shaken in the same conditions and sampled periodically for measurement of OD600. Growth rate of each culture was determined by non-linear regression using an exponential growth equation (KaleidaGraph v5) and values obtained from replicates were averaged.

### RT-PCR experiments

Strains were grown overnight at 30°C to mid-log phase in SCD medium. For each sample, cells were diluted to OD600 = 0.2 in fresh SCD medium with or without the stressor and shaken at 30°C. At the appropriate time points, 1.5 mL of the culture was harvested by centrifugation (14 k, 30 sec) and the supernatant aspirated. Cell pellets were flash frozen in liquid nitrogen and stored at −80°C until RNA extraction. Total RNA was extracted from cells using a hot acid phenol-chloroform extraction protocol (73). Briefly, cells were lysed in an RNA lysis buffer (NaOAc, EDTA, SDS) and then RNA was purified through two phenol extractions and a final chloroform extraction. RNA was precipitated with isopropanol and resuspended in TE buffer. RNA extracts were treated with DNAse (New England Biolabs) to ensure no genomic DNA contamination. 500 ng of RNA was reverse transcribed using the High-Capacity cDNA Reverse Transcription Kit (Thermo). Real-time PCR was performed using the CFX96 Touch Real-Time PCR Detection System (BioRad) using the ABsolute Blue QPCR Mix, SYBR Green Kit (ThermoFisher) with the following parameters: 95°C 15:00 Min, 40x (95°C 00:15, 58°C 00:30, 72°C 00:30). *CDR1* transcript levels were normalized to *TEF1* transcript levels in each sample and this ratio from all samples were normalized to that of untreated BG14 cells.

### Western blots

Cells were grown in SCD-ura medium to OD600 = 0.5 then harvested and extracted with 10% TCA. Pellets were suspended in SDS sample buffer containing 8M urea, heated to 80°C, then separated on 10% SDS gels and transferred to PVDF membranes. The membranes were blocked with 10% nonfat milk, probed sequentially with antibodies specific for Erg11 (45) or Pgk1 (Invitrogen) and visualized on a BioRad imager using HRP-conjugated secondary antibodies (Jackson ImmunoResearch) and Western Blotting Detection Reagents (GE).

### Competitive growth assay

Colonies of BG14 and *cyb5Δ* mutants were grown overnight to saturation, mixed at 1:10 ratios, then diluted 1:100 into fresh SCD media supplemented with 128 µg/mL of fluconazole. Cells were allowed to grow overnight at 30°C, pelleted, washed, and resuspended in fresh media lacking fluconazole and grown overnight. Aliquots were removed after mixing and after regrowth, diluted, and plated on YPD plates for 2 days at 30°C. CFUs were determined before and after replica plating onto SCD-Ura plates.

## ACKNOWLEDGMENTS

The authors thank Drs. Scott Moye-Rowley, Alejandro de las Peñas, and Taiga Miyazaki for generously providing antibodies, *C. glabrata* strains, and advice. We thank Drs. Winston Timp and John Kim for providing access to critical instruments. We thank Abigail Harrington and Joshua Schultz for thoughtful comments on the manuscript. This research was supported by grants from the National Institutes of Health (T32-GM007231 to the JHU CMDB training program; R01-AI153414 to KWC; R01-AI046223 to BPC).

---

**Supplemental Figure 1.**
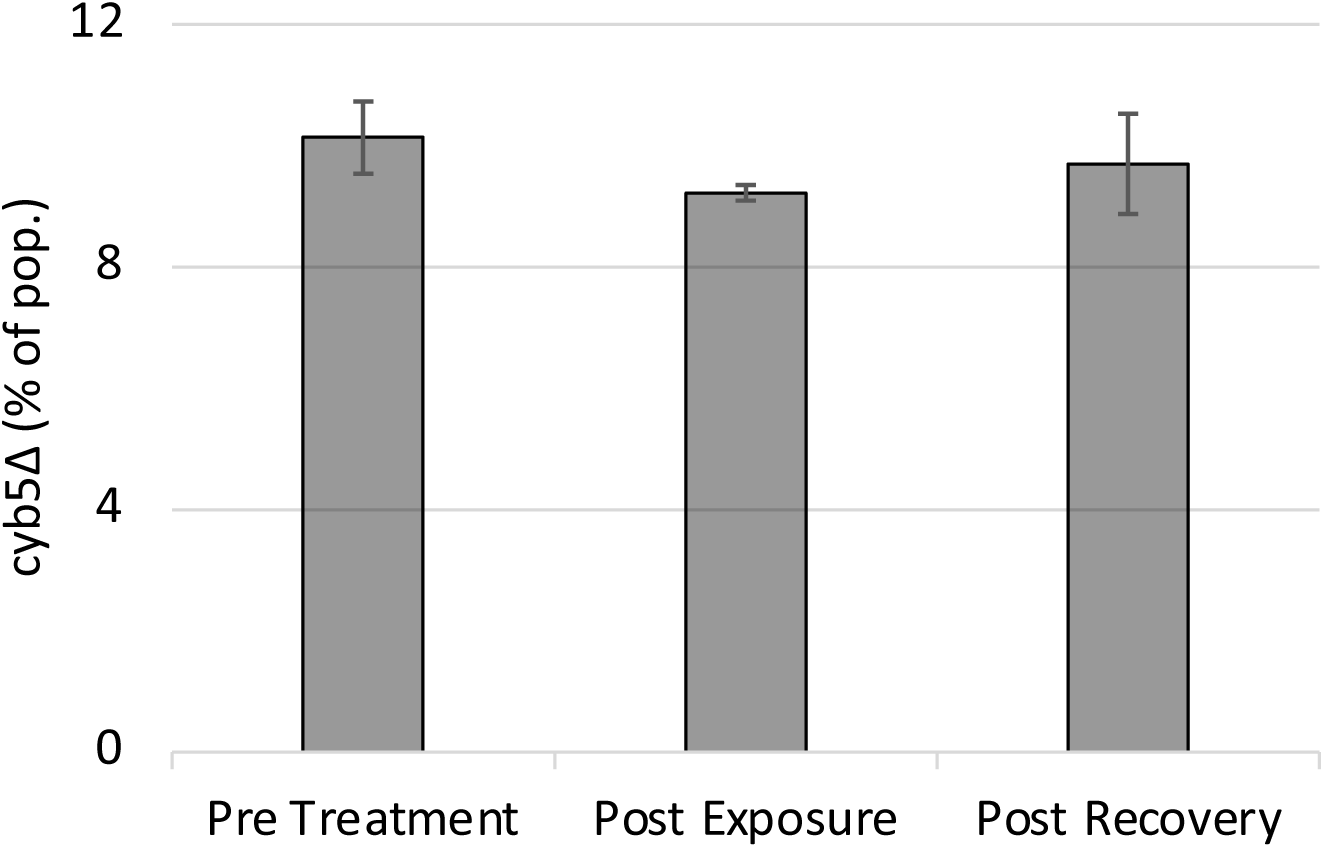
Fluconazole sensitivity of *cyb5Δ* is rescued by co-culturing with BG14. Wild-type and *cyb5Δ* cells were grown to saturation and mixed at a 1:10 *cyb5Δ* to wild-type ratio. Mixtures were back diluted 1:100 into SCD media and exposed to 128 µg/mL of fluconazole for 24 hr. Cells were washed and resuspended in fresh SCD media for 24 hours. Relative abundance of wild-type to *cyb5Δ* mutants was calculated at the following time points, before drug exposure, post drug exposure and after recovery. Values are averaged from 3 biological replicates (±SD).

**Supplemental Figure 2.**
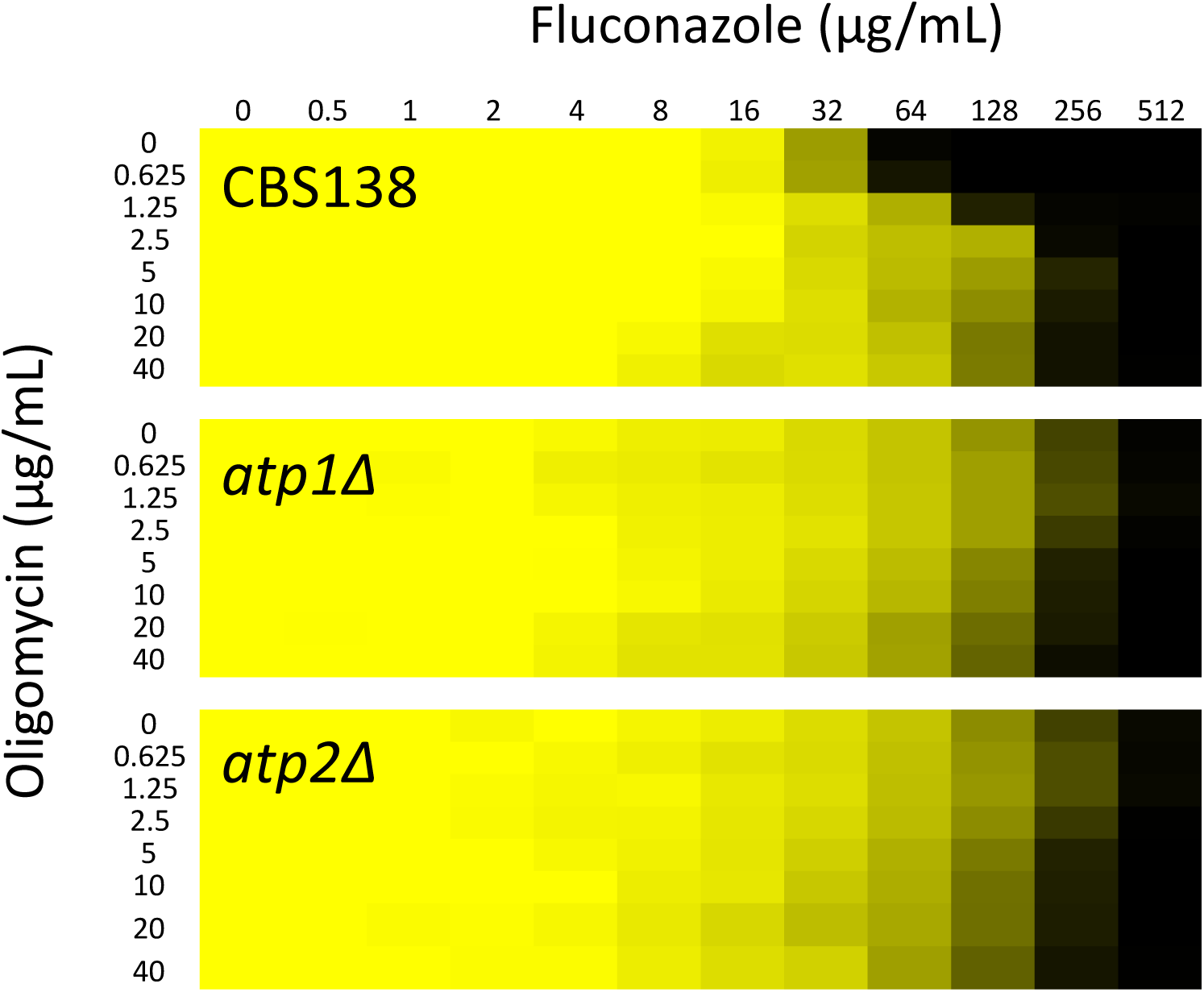
Oligomycin antagonism of fluconazole effectiveness depends on ATP synthase. Growth of the wild-type CBS138, *atp1Δ,* and *atp2Δ* strains was measured at 600 nm after 20 hr incubation in SCD medium containing varying concentrations of fluconazole plus varying concentrations of oligomycin. Data are plotted as a heat map from saturation (yellow) to no growth (black).

**Supplemental Figure 3.**
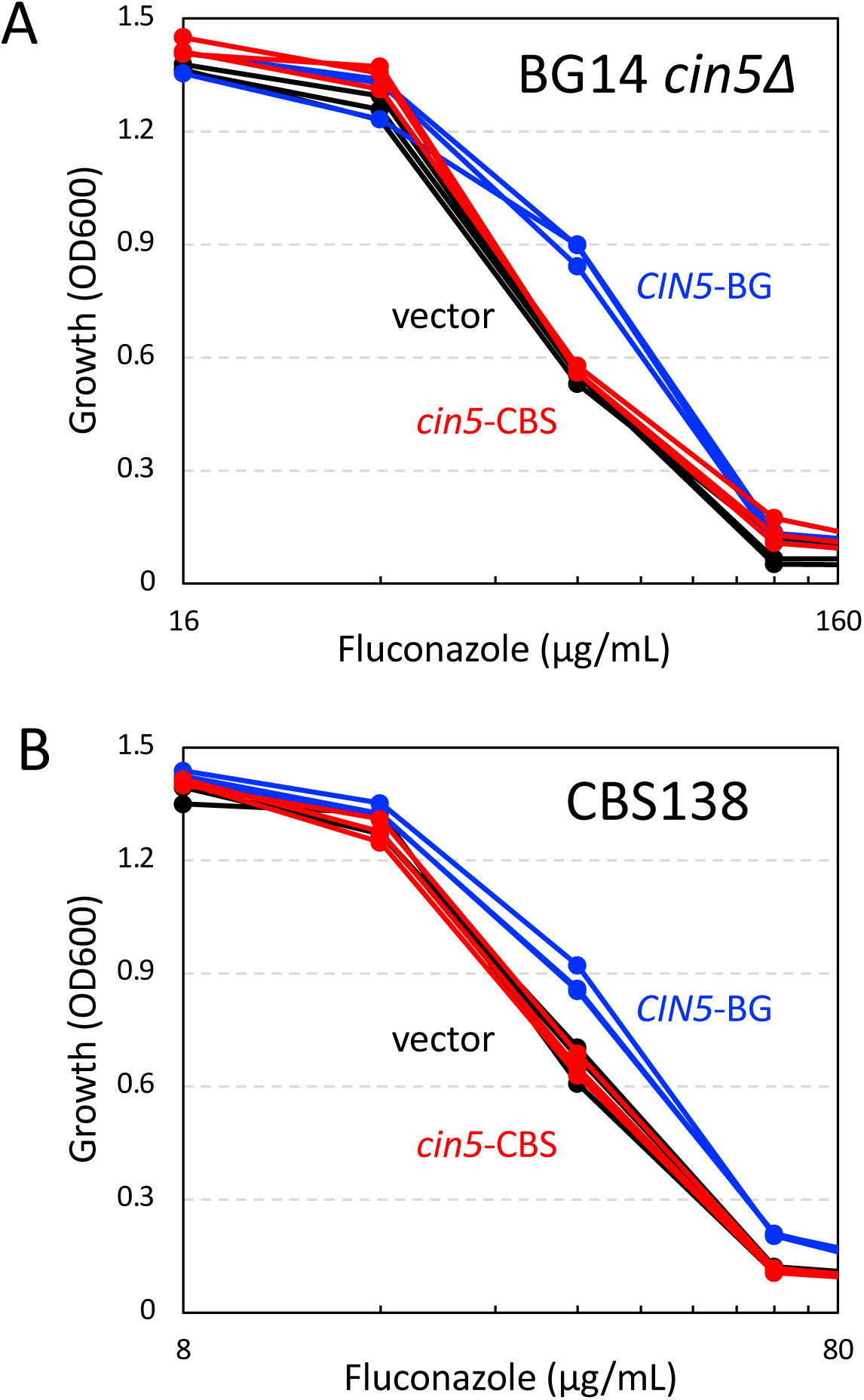
An early stop codon in CBS138 inactivates *CIN5* function. Plasmids bearing *CIN5* from either BG2 or CBS138 parent strains were transformed into the BG14 *cin5Δ* (A) and CBS138 (B) strains and assayed in triplicate for susceptibility to fluconazole.

**Supplemental Figure 4.**
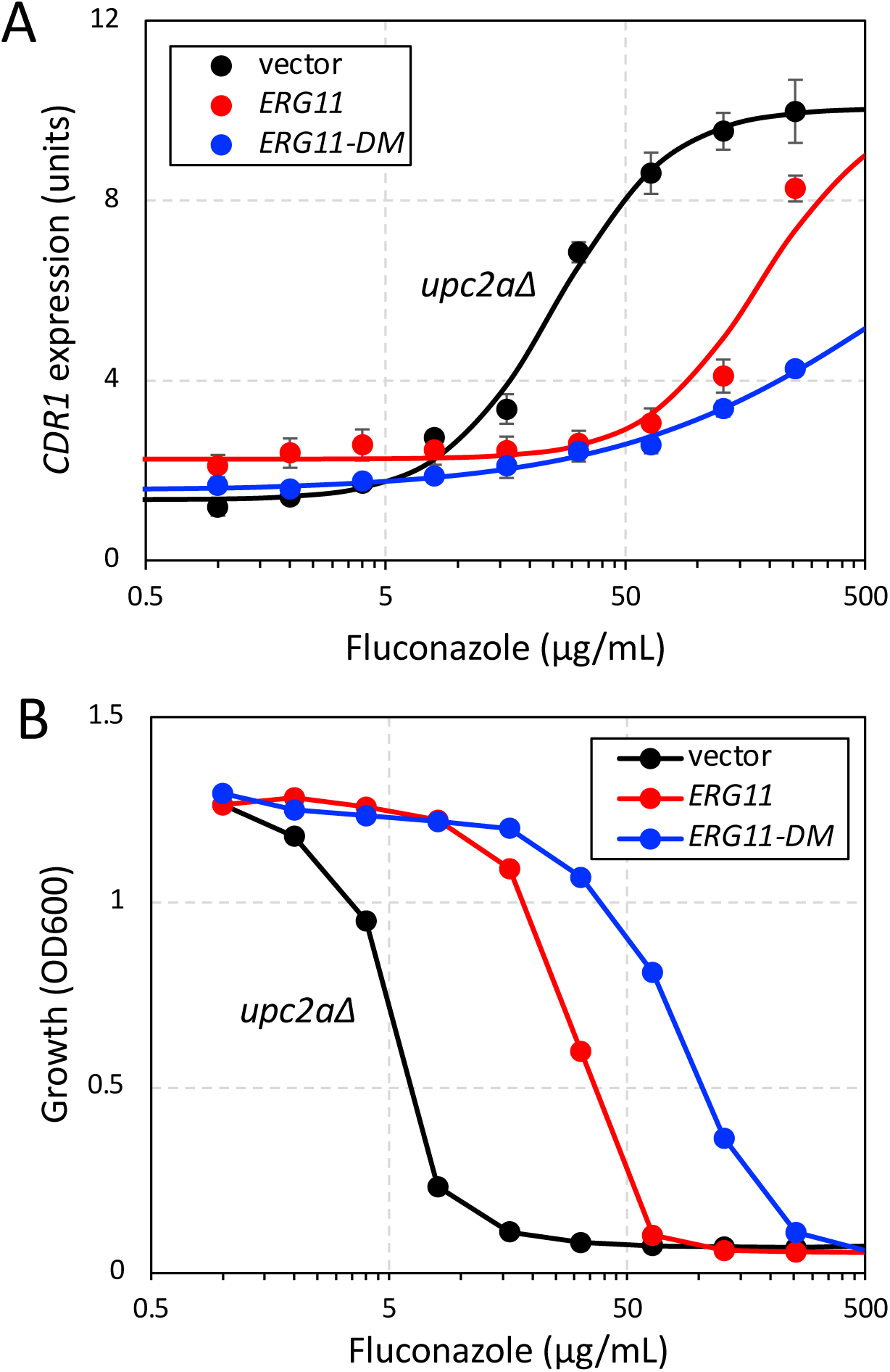
Overexpressed Erg11 and Erg11-DM confer fluconazole resistance independent of Upc2A. The *upc2aΔ* mutant transformed with empty vector, *ERG11*, and *ERG11-DM* overexpression plasmids were grown in SCD-ura medium containing the indicated levels of fluconazole and analyzed for *CDR1* expression (A) and growth (B) as described in Fig. 6.

